# Heterologous protection against Asian Zika virus challenge in rhesus macaques

**DOI:** 10.1101/059592

**Authors:** Matthew T. Aliota, Dawn M. Dudley, Christina M. Newman, Emma L. Mohr, Dane D. Gellerup, Meghan E. Breitbach, Connor R. Buechler, Mustafa N. Rasheed, Mariel S. Mohns, Andrea M. Weiler, Gabrielle L. Barry, Kim L. Weisgrau, Josh A. Eudailey, Eva G. Rakasz, Logan J. Vosler, Jennifer Post, Saverio Capuano, Thaddeus G. Golos, M. Anthony Moody, Sallie R. Permar, Jorge E. Osorio, Thomas C. Friedrich, Shelby L. O’Connor, David H. O’Connor

**Author notes:** Address correspondence to David H. O’Connor. These authors contributed equally to this work (Matthew T. Aliota, Dawn M. Dudley).

## Abstract

Zika virus (ZIKV) isolates are genetically diverse, but belong to two recognized lineages, termed “African” and “Asian.” Asian ZIKV infection during pregnancy causes fetal abnormalities including microcephaly. Developing an effective preventative Zika virus vaccine that protects pregnant women is essential for minimizing fetal abnormalities; at least 18 groups are developing ZIKV vaccines (Hayden, 2016). The genetic and antigenic variability of many RNA viruses limits the effectiveness of vaccines, and the degree to which immunity against one ZIKV strain could provide protection against another is unknown. Here we show that rhesus macaques infected with East African ZIKV strain MR766 are completely protected from subsequent infection with heterologous Asian ZIKV. MR766 is more genetically divergent from all known Asian ZIKV strains than Asian ZIKV strains are from one another. Therefore, ZIKV strain selection is unlikely to compromise vaccine effectiveness.

**Highlights:**

- African Zika virus (ZIKV) strain MR766 productively infects macaques (68 characters)
- Immunity elicited by MR766 protects macaques against heterologous Asian ZIKV (77 characters)
- In vivo restoration of a putative N-linked glycosylation site in MR766 (70 characters)
- Immunogen selection is unlikely to adversely affect the breadth of vaccine protection (85 characters)

**eTOC:** An effective Zika virus vaccine is needed to prevent infection-associated fetal abnormalities. Macaques whose immune responses are primed by infection with East African ZIKV are completely protected from reinfection with heterologous Asian ZIKV. Any Asian ZIKV immunogen that protects against homologous challenge will likely confer protection against all other Asian ZIKV strains.

## Introduction

Zika virus (ZIKV) is an arthropod-borne member of the genus *Flavivirus* of the Spondweni serocomplex that is currently causing an explosive outbreak of febrile disease in the Americas. Historically, ZIKV existed in relative obscurity with only sporadic confirmed human infections until the end of the last century (Hayes, 2009). ZIKV is believed to have originated in Africa, where it is maintained in an enzootic cycle that includes unknown vertebrate hosts (nonhuman primates are suspected) and arboreal Aedes mosquitoes (HADDOW et al., 1964; McCrae and Kirya, 1982; Wolfe et al., 2001). In fact, ZIKV was first isolated from the blood of a sentinel rhesus monkey during yellow fever virus surveillance studies in the Zika forest of Uganda (DICK et al., 1952). The virus is thought to have spread from East Africa into both West Africa and Asia ~50-100 years ago (Faye et al., 2014). Beginning in 2007, ZIKV outbreaks were reported in Yap Island of the Federated states of Micronesia (Duffy et al., 2009), French Polynesia (Cao-Lormeau et al., 2014), other Pacific islands (Chang et al., 2016), and in early 2015, in the state of Rio Grande do Norte in northern Brazil (Zanluca et al., 2015). Since its introduction into the Americas, ZIKV has spread essentially uncontrolled with at least 39 countries and territories experiencing autochthonous transmission, including the US territory of Puerto Rico (Pan American Health Organization, 2016). Eventually, local spread in the southern United States seems likely. In humans, ZIKV infection typically causes a mild and self-limiting illness known as Zika fever, which often is accompanied by maculopapular rash, headache, and myalgia (Campos et al., 2015; Camacho et al., 2016). During the current outbreak, a causal relationship between prenatal ZIKV infection and fetal microcephaly, as well as other serious brain anomalies, has been established (Driggers et al., 2016; Mlakar et al., 2016; Rasmussen et al., 2016). Development and testing of vaccines that elicit protective immune responses among girls and women before pregnancy is a top public health priority (Cohen, 2016).

The ZIKV genome is an ~11 kb single-stranded, positive sense RNA that contains a single open reading frame. Once the RNA genome is released into the cytoplasm it is directly translated into a polyprotein precursor. The polyprotein is subsequently glycosylated by cellular glycosyltransferases and cleaved by a combination of viral and host proteases to release three structural (C, prM, and E) and seven nonstructural proteins (NS1, NS2a, NS2b, NS3, NS4a, NS4b, and NS5) (Sirohi et al., 2016). The envelope (E) glycoprotein is a target for broadly protective neutralizing antibodies in ZIKV and other flaviviruses and is an attractive candidate immunogen for inclusion in ZIKV vaccines (Dai et al., 2016). Understanding the breadth of immunity elicited by the envelope glycoprotein and the host selection of viral variants is therefore important for vaccine design.

There are three distinct lineages of ZIKV: West African (Nigerian cluster), East African (MR766 prototype cluster), and Asian (Lanciotti et al., 2016). All of the ZIKV strains circulating in the Western hemisphere are Asian lineage. The E amino acid identity among all Asian lineage ZIKVs is >99%, and as a group, these are only ~96% and 97% amino acid identical to representative East African and West African viruses, respectively (Lanciotti et al., 2016). It is not known whether the differences between African and Asian lineage ZIKV have any phenotypic impact, e.g., increased transmissibility or pathogenicity. Because human infections with ZIKV have historically been sporadic, and, until recently, limited to small-scale epidemics, neither the disease caused by ZIKV nor the molecular determinants of immunity have been well characterized. Accordingly, we recently developed an animal model for Asian-lineage ZIKV infection in Indian-origin rhesus macaques (Macaca mulatta)(Dudley et al., 2016), and demonstrated that immune responses elicited by infection with Asian ZIKV completely protect against homologous rechallenge with the same ZIKV strain. While this demonstrated the potency of naturally elicited antiviral immunity, it did not address whether such immunity is broadly protective against heterologous ZIKV strains.

To investigate the breadth of protective ZIKV immunity between heterologous lineages of the virus, we infected three macaques with the African prototype strain of ZIKV, MR766 (DICK et al., 1952). All three animals exhibited an acute, self-limiting infection similar to those previously observed in macaques infected with Asian ZIKV. Immune responses against MR766 completely protected all three macaques against reinfection using an Asian ZIKV strain and dose that productively infected 6/6 naïve macaques.

## Results

### Sequence characterization of African lineage MR766 isolates

We elected to use the African ZIKV strain MR766 as the primary challenge virus since this is the prototypical East African ZIKV used in previous studies. MR766 was derived from the original isolate from the Zika Forest, Uganda (DICK et al., 1952). Interestingly, GenBank contains records for seven different sequences all called ZIKV prototype strain “MR766” (accession numbers: DQ859059, AY632535, LC002520, KU963573, KU955594, KU720415, HQ234498). Differences among these MR766 sequences have been noted previously (Haddow et al., 2012), but not extensively characterized. All of the Genbank sequences are 99.7-100.0% nucleotide identical to one another within the polyprotein coding sequence, with the exception of DQ859059. Others have shown that the sequence of DQ859059 matches a mosquito-derived sequence unrelated to MR766 (Loman, 2016). This sequence should be considered a database error and not used as the prototype for any future MR766 analyses.

The most obvious difference between the Genbank MR766 sequences is a four amino acid sequence in the ‘150 loop’ of the E protein that contains a potential N-linked glycosylation site (Dai et al., 2016) that is absent from some of the GenBank reference sequences. The ‘150 loop’ is also absent from other flaviviruses (Dai et al., 2016). We explored this deletion in greater detail by deep sequencing three MR766 isolates, Zika virus/R.macaque-tc/UGA/1947/MR766-3329 (hereafter referred to as challenge stock, or abbreviated Chal Stk in figures), WRCEVA, and CDC (see Table 1 for passage history). Figure 1A shows amino acid sites in E that differ between the MR766 Genbank sequences and deep sequenced MR766 strains. The deletion was present in between 80.0% and 100.0% of sequencing reads from the three deep sequenced isolates. 85.7% of reads in the stock used to infect the animals in this study (i.e., ‘Chal Stk’ in Figure 1A) contained the deletion. Neither of the other sites predicted to have amino acid variability among Genbank MR766 isolates in E differ in any of the three isolates examined in this study.

**Table 1.** Deep sequenced MR766 isolates. WRCEVA= World Reference Center for Emerging Viruses and Arboviruses at the University of Texas Medical Branch, CDC= Centers for Disease control and Prevention, SM= suckling mouse, Vero= African green monkey kidney cells, C6/36= Aedes albopictus cells.

| Stock Name | # of SM Passage | # Vero cell Passage | #C6/36 cell Passage |
| --- | --- | --- | --- |
| ZIKV MR766 WRCEVA | 150 | 1 | 1 |
| ZIKV MR766 CDC | 149 | 2 | 0 |
| ZIKV MR766 (challenge stock) | 149 | 2 | 1 |

Including the in-frame four-amino-acid deletion, MR766 is ~96% amino acid identical in E to Asian ZIKV isolates (Figure 1B). In contrast, Asian ZIKV isolates are more than 99% amino acid identical to one another in envelope, typically differing by only 0-2 amino acids. In other words, the MR766 challenge stock is more genetically divergent from Asian ZIKV than Asian ZIKV strains are from one another. Therefore, we hypothesized that if immunity elicited by infection of macaques with MR766 protects against reinfection with Asian ZIKV infection, immunity elicited by any Asian ZIKV infection should be sufficient to confer complete protection against subsequent Asian ZIKV reinfection.

### Primary infection of Indian-origin rhesus macaques with East African Zika virus MR766

To examine the course of primary infection with MR766, we infected Indian-origin rhesus macaques (two males and one female). This group of animals was designated ZIKV-002 to provide consistency with real-time data on these animals that is publicly available at (Zika experimental science team, 2016). Animals were inoculated subcutaneously with either 1×10^4^, 1×10^5^, or 1×10^6^ PFU/mL of our challenge stock, consistent with the route and challenge doses of two previously published cohorts (ZIKV-001 and ZIKV-004) of Indian-origin rhesus macaques challenged subcutaneously with Zika virus/H.sapiens-tc/FRA/2013/FrenchPolynesia-01_v1c1, an Asian ZIKV termed ‘ZIKV-FP’ for the remainder of the manuscript (Dudley et al., 2016). A study schematic is shown in Figure 2A. All three animals were productively infected with MR766, with detectable plasma viremia one day post inoculation (dpi) in two of the three animals and in all three by two dpi using a qRT-PCR that amplifies MR766 and ZIKV-FP with essentially identical efficiency (Figure S1). Plasma viremia peaked in all animals between two and five dpi, and ranged from 2.21 × 10^4^ to 2.64 × 10^5^ vRNA copies/mL. The highest plasma viremia was observed for the animal inoculated with the lowest primary challenge dose (1 × 10^4^ PFU/mL); the peak of plasma viremia also occurred later in this animal (five dpi) than in the other two. By ten dpi, plasma viral loads were undetectable in all three animals. Cerebrospinal fluid (CSF) was sampled at four dpi and 14 dpi, and was positive for vRNA in animal 295022 on day four (955 vRNA copies/mL) and 405734 on day 14 post-infection (937 vRNA copies/mL). 562876 was negative at both CSF collection timepoints. Detection of ZIKV RNA in other body fluids (saliva and urine) generally lagged behind detection in plasma by two to seven days. Viral RNA was detected in the saliva of two animals by seven dpi, in the third animal by nine dpi, and ranged from 3.8 × 10^1^ to 2.6 × 10^4^ vRNA copies/ mL (Figure 2C). Viral RNA was detected in passively collected pan urine from only 295022 (Figure 2D). After 14 dpi, no animals had detectable vRNA in any body fluids at the remainder of the sampled timepoints.

**Figure 1.**
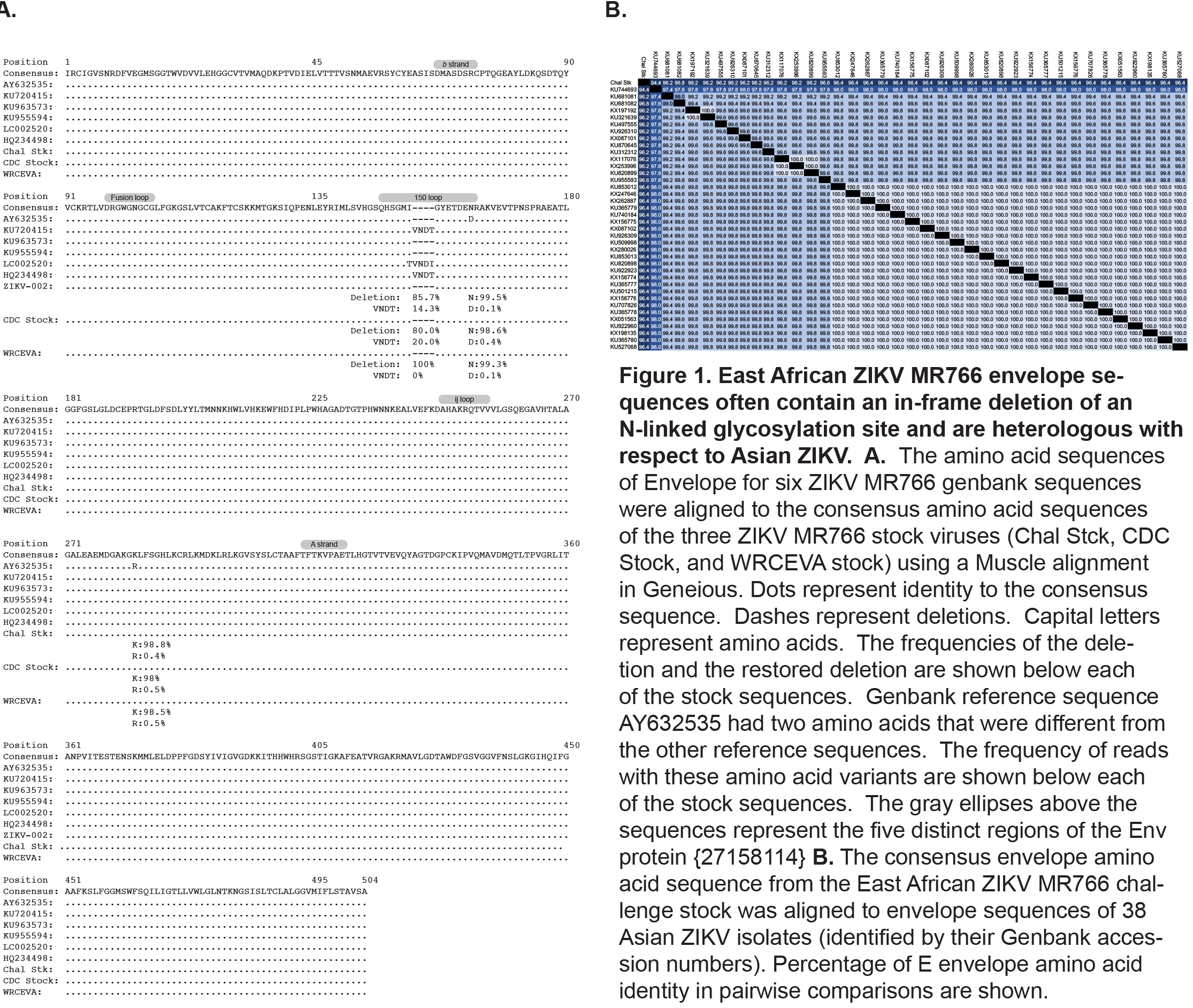
East African ZIKV MR766 envelope sequences often contain an in-frame deletion of an N-linked glycosylation site and are heterologous with respect to Asian ZIKV. A. The amino acid sequences of Envelope for six ZIKV MR766 genbank sequences were aligned to the consensus amino acid sequences of the three ZIKV MR766 stock viruses (Chal Stck, CDC Stock, and WRCEVA stock) using a Muscle alignment in Geneious. Dots represent identity to the consensus sequence. Dashes represent deletions. Capital letters represent amino acids. The frequencies of the deletion and the restored deletion are shown below each of the stock sequences. Genbank reference sequence AY632535 had two amino acids that were different from the other reference sequences. The frequency of reads with these amino acid variants are shown below each of the stock sequences. The gray ellipses above the sequences represent the five distinct regions of the Env protein {27158114} B. The consensus envelope amino acid s equence from the East African ZIKV MR766 challenge stock was aligned to envelope sequences of 38 Asian ZIKV isolates (identified by their Genbank accession numbers). Percentage of E envelope amino acid identity in pairwise comparisons are shown.

**Figure 2.**
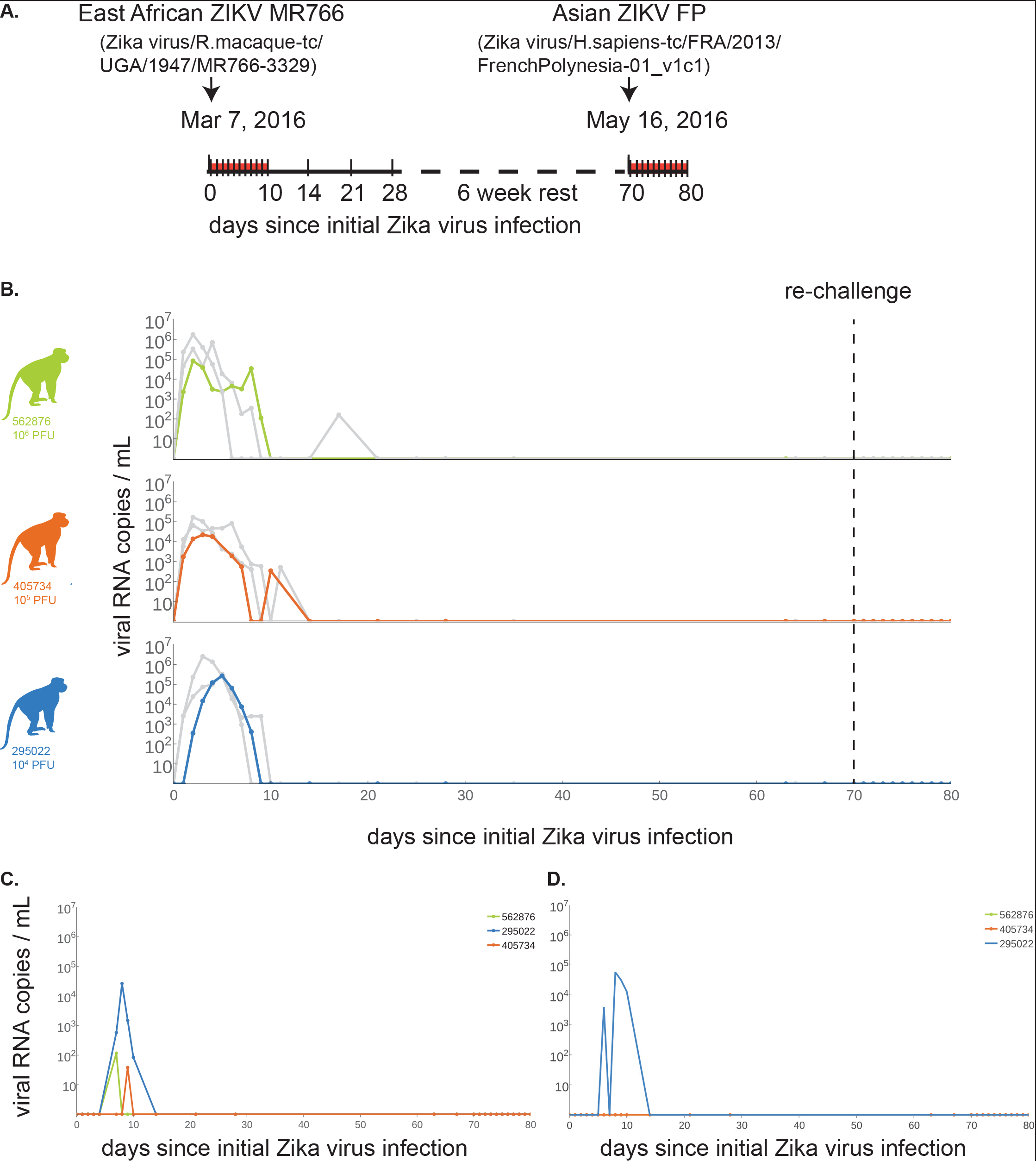
ZIKV-002 macaques challenged with East African ZIKV MR766 are protected from heterologous reinfection with Asian ZIKV FP. A. Study timeline with dates of primary and secondary heterologous ZIKV challenges. Samples were collected daily from 0 to 10 dpi, and then weekly thereafter until secondary challenge (denoted by ticks along the timeline). Challenge stocks were derived from the East African and French Polynesian virus strains detailed above the timeline. B. Plasma viral loads, shown as vRNA copies/mL for each of the macaques challenged with 1 ×10^6^ (solid green line), 1× 10^5^ (solid orange line), or 1 × 10^4^ (solid blue line) PFU/mL of ZIKV MR766 from the date of primary challenge through 10 days post heterologous challenge with Asian ZIKV. For comparison of plasma viral loads between ZIKV strains, solid light grey lines depict the plasma viral load trajectories for animals that were challenged with the same dose of Asian ZIKV FP and then rechallenged with homologous Asian ZIKV FP. C. Oral swab and D. pan urine viral loads.

### In-frame envelope deletion in MR766 stocks is rapidly lost in vivo

Three and six days after infection, vRNA from the animals was reverse transcribed and PCR-amplified using a primer pair that amplifies the E coding region, including the 12nt in-frame deletion in the 150 loop. PCR amplicons were randomly fragmented and deep sequenced. The in-frame deletion was detected in no more than 0.4% of all reads by three days post infection (Figure 3). This suggests that the minority population in the challenge stock containing an intact 150 loop rapidly outcompeted viruses containing the in-frame deletion. Because the sequence of the intact sequence was identical to the sequence of the minority population of the challenge stock, de novo repair of the deletion by three days post-infection is unlikely.

### Robust cellular and humoral immunity to ZIKV

Proliferation of CD8+ and CD4+ T cells, as well as natural killer cells, was observed following ZIKV MR766 challenge (Figure 4A-C). These responses, peaking six to ten days post challenge, were robust and mirrored the kinetics of proliferation we previously observed in macaques infected with ZIKV-FP. The peak magnitude of proliferation was slightly lower, on average, than observed following ZIKV-FP infection but the small number of macaques in the study makes it impossible to quantify the significance of this observation. Similarly, we detected an increase in the number of antibody-producing plasmablasts in all three animals after MR766 infection (Figure 4D). Serum neutralizing antibody responses also were measured by plaque reduction neutralization test (PRNT90), and all animals exhibited neutralizing antibody (nAb) titers ≥20 against both the challenge stock and ZIKV FP seven days prior to rechallenge (Figure 4E).

### Heterologous challenge with Asian ZIKV

To determine if primary infection with East African ZIKV results in protection from heterologous re-challenge with Asian ZIKV, we inoculated the ZIKV-002 animals with 1 × 10^4^ PFU/mL of ZIKV-FP at 70 dpi (10 weeks after primary challenge). This dose was chosen based on successful infection of 6/6 animals from multiple cohorts challenged with this dose to date ((Dudley et al., 2016) and zika.labkey.com). Viral RNA was undetectable in plasma (Figure 2B), saliva (Figure 2C), and urine (Figure 2D) at all timepoints through 10 days post re-challenge.

## Discussion

Here we describe the first evidence demonstrating that protective immunity resulting from natural ZIKV infection confers protection against reinfection with a heterologous strain of the virus. These findings have potential implications for vaccine development and implementation. Although ZIKV immunology is still in its infancy, we hypothesize that, similar to DENV, nAbs are a critical component of the protective immune response (Guzman et al., 2007; SABIN, 1952). However, unlike DENV where individuals are at increased risk of symptomatic infection and severe disease upon subsequent heterologous infection, primary ZIKV infection was protective against heterologous rechallenge (though DENV serotypes are more genetically dissimilar to each other than East African and Asian ZIKV). It should be noted that with DENV, cross-serotype protection against symptomatic infection has been observed for up to two years after primary infection, after which point individuals were at greater risk of severe disease (Anderson et al., 2014; Reich et al., 2013; Montoya et al., 2013), because cross-serotype-reactive antibodies are believed to decay to sub-neutralizing levels that bind DENV without neutralization. Therefore, further work is needed (both experimental and epidemiological) to understand the duration of protective ZIKV immunity after natural infection. Moreover, our study is limited because re-challenges were performed at only a single timepoint; we do not know how soon after primary infection protective immune responses emerge, or for how long they might endure.

**Figure 3.**
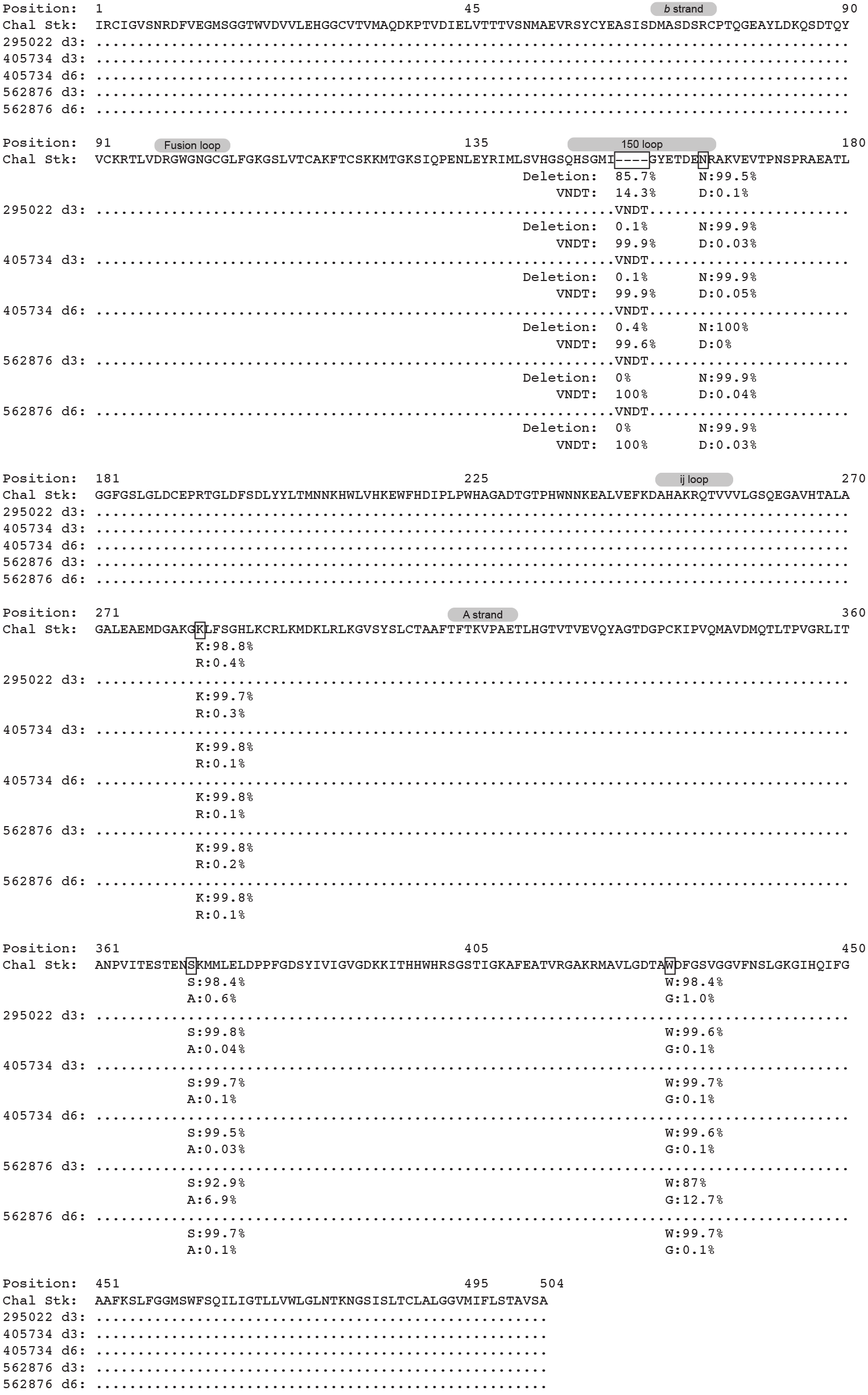
An N-linked glycosylation site in envelope is rapidly selected in vivo. Envelope sequences from the three animals were sequenced at three days post infection, and from two of the animals at day six post infection. A MUSCLE alignment of the translated sequences was generated in Geneious. Dots represent identity to the consensus sequence. Dashes represent deletions. Capital letters represent amino acids. The frequencies of the deletion and the restored deletion are shown below each of the stock sequences, with the indicated site boxed. Amino acid variant frequencies matching the variant sites in Figure 1A are shown. There were two additional nonsynonymous variants at greater than 5% in animal 562876 at day 3, and the frequency of the amino acid variants from the other animals and time points are shown below each sample. The gray ellipses above the sequences represent the five distinct regions of the Env protein {27158114}.

**Figure 4.**
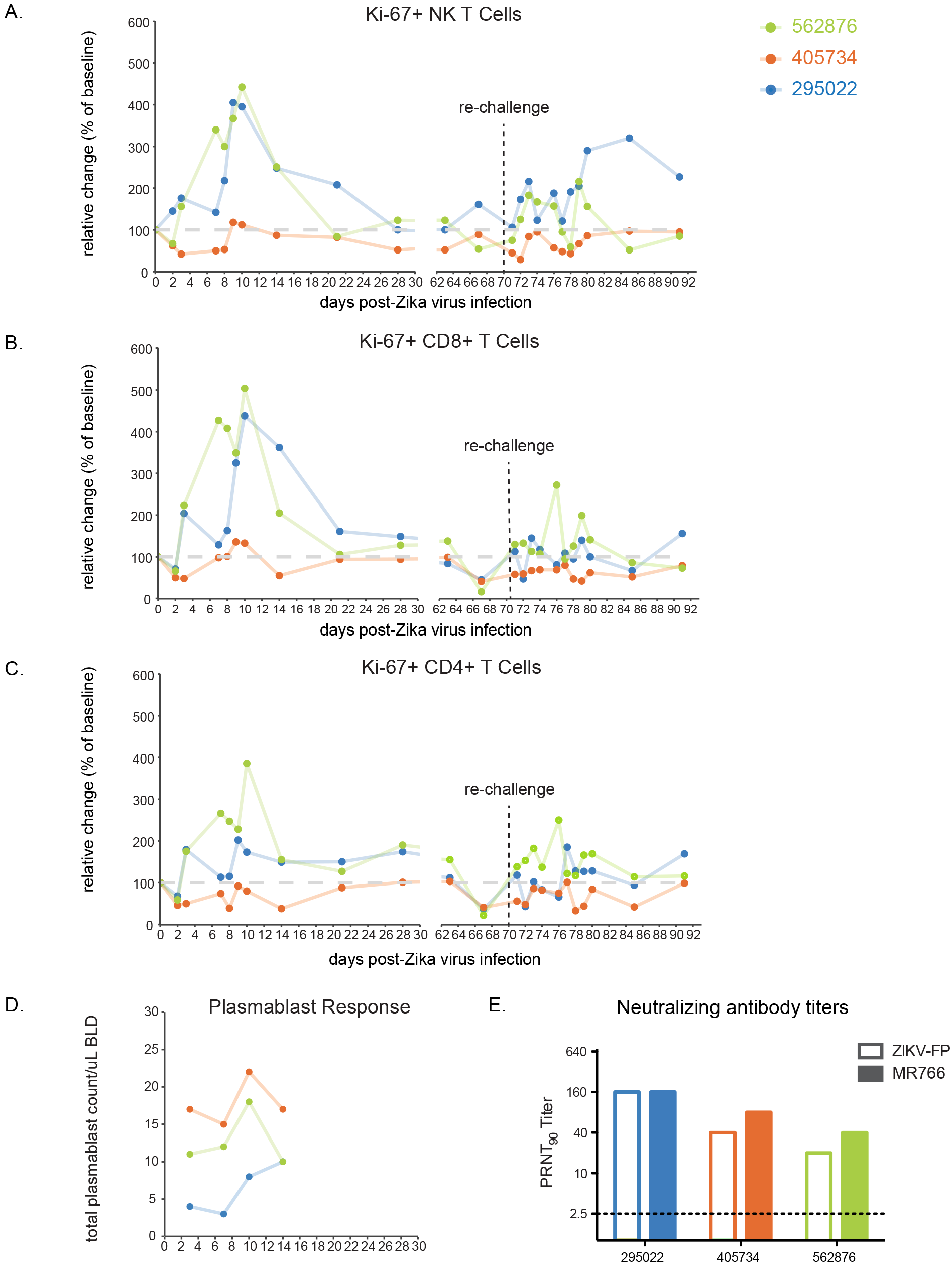
East African ZIKV MR766infectionelicits a robust, multifacetedimmuneresponse. Expansion of Ki-67+ (activated) A. NK cells, B. CD8+ T cells, and C. CD4+ T cells was measured at days 0, 2, 3, and days 7 through 10, then on days 14, 21 and 28 post-infection. After a rest period, activated immune responses were measured at days 63 and 67 post-infection. At 70 dpi, animals were heterologously re-challenged with Asian ZIKV FP. Expansion of activated cells was measured daily through 80 dpi, then at days 85 and 91 post-infection. Absolute numbers of activated cells/μl of blood are presented relative to the baseline value set to 100%. D. Total number of plasmablast cells were found in PBMCs collected at days 3, 7, 10 and 14 post-infection for each animal. E. PRNT90 titers seven days prior to rechallenge for ZIKV002 animals against Asian ZIKV FP (open bars) and East African ZIKV MR766 (filled bars).

Incidental to the primary observation of protective heterologous ZIKV immunity, we also demonstrated that selection appears to favor maintenance of four amino acids that are deleted in the majority of viral sequences in the challenge stock. It is interesting to note that this region contains a putative N-linked glycosylation site (E_154_), leading us to speculate that this might be consequential for in vivo replication. Flaviviruses contain several putative N-linked glycosylation sites (N-X-S/T) in the E and NS1 proteins. The glycosylation pattern on the viral envelope protein varies among flaviviruses and even among strains of the same virus (Kaufmann and Rossmann, 2011). Some, but not all, African ZIKV strains contain a putative E glycosylation site at E_154_ (Haddow et al., 2012). This is also true of West Nile virus (WNV) E_154_. N-linked glycosylation plays an important role in both the assembly and the infectivity of many viruses (Courageot et al., 2000; Lin et al., 2003; Schlesinger et al., 1985; Tassaneetrithep et al., 2003). For WNV, E protein glycosylation can influence virus infectivity (Hanna et al., 2005). Deglycosylation of both E and NS1 proteins of WNV completely attenuated neuroinvasiveness and induced protective immunity in the murine model with low doses of virus (Whiteman et al., 2010). It has been hypothesized that extensive mouse brain or cell culture passage could lead to the deletion of the potential glycosylation site (Chambers et al., 1998); therefore it is important to note that the African ZIKV strains analyzed here all underwent extensive mouse brain passage. Consequently, it will be important to sequence low passage, geographically distinct strains to confirm whether or not this glycosylation site polymorphism is an artifact of passage history or if it is representative of circulating strains. However, the fact that the deletion was likely selected against in vivo supports the hypothesis that passage history has influenced glycosylation sites in the African prototype strain and suggests that preservation of this glycosylation site may be important for efficient replication in primates.

These results have important implications for vaccine design and testing. MR766 is more genetically dissimilar to Asian lineage ZIKV isolates than any two Asian lineage ZIKV isolates are from one another. Unlike vaccines for other RNA viruses where immunogen selection is critical, our results suggest that protective immunity elicited against any Asian ZIKV should be sufficient to confer broad protection against all Asian ZIKV strains. Together with our previous results, demonstrating that previous Asian-lineage ZIKV infection protects macaques from homologous rechallenge, the results shown here suggest that immunity elicited by a single ZIKV antigen may provide cross-protective immunity against a multitude of ZIKV strains.

The complete protection from homologous and heterologous rechallenge also suggests that the immunologic barrier for complete protection may be comparatively low, such that vaccines with acceptable safety may have desirable efficacy even if they are not highly immunogenic. We do not currently know the correlate of protective immunity in these animals, since robust T cell, NK cell, B cell, and nAb responses were elicited, but subsequent work with individual lymphocyte subsets in cell depletion or adoptive transfer experiments, as well as passive antibody transfer experiments, should clarify this important issue.

## Experimental procedures

**Study Design**. This study was designed to examine whether prior infection with African Zika virus (ZIKV) provides protection from heterologous challenge with an Asian ZIKV isolate in the rhesus macaque model. Datasets used in this manuscript are publicly available from http://go.wisc.edu/50bfn2

**Ethical approval**. This study was approved by the University of Wisconsin-Madison Institutional Animal Care and Use Committee (Animal Care and Use Protocol Number G005401).

**Animals**. Two male and one female, Indian-origin rhesus macaques (Macaca mulatta) utilized in this study were cared for by the staff at the Wisconsin National Primate Research Center (WNPRC) in accordance with the regulations, guidelines, and recommendations outlined in the Animal Welfare Act, the Guide for the Care and Use of Laboratory Animals, and the Weatherall report. In addition, all macaques utilized in the study were free of Macacine herpesvirus 1, Simian Retrovirus Type D, Simian T-lymphotropic virus Type 1, and Simian Immunodeficiency Virus. For all procedures, animals were anesthetized with an intramuscular dose of ketamine (10mL/kg). Blood samples were obtained using a vacutainer or needle and syringe from the femoral or saphenous vein. Macaques challenged with Asian ZIKV (Zika virus/H.sapiens-tc/FRA/2013/FrenchPolynesia-01_v1c1; ZIKV FP) were used for comparison; details on these animals are available in (Dudley et al., 2016).

**Viruses**. ZIKV prototype strain MR766 (referred to as CDC) was obtained from Brandy Russell (CDC, Ft. Collins, CO) and was originally isolated from a sentinel rhesus monkey in 1947 from the Zika Forest, Entebbe, Uganda and passaged 149 times through suckling mouse brains and twice on Vero cells. ZIKV French Polynesian strain (ZIKV FP) was obtained from Xavier de Lamballerie (European Virus Archive, Marseille, France). It was originally isolated from a 51-year old female in France after travel to French Polynesia in 2013 and passaged a single time on Vero cells. Challenge virus stocks were prepared by inoculation onto a confluent monolayer of C6/36 mosquito cells. A single, clarified harvest of each virus, with titers of 5.9 × 10^6^ PFU/mL (3.9 × 10^9^ vRNA copies/mL) and 1.26 × 10^6^ PFU/mL (1.43 × 10^9^ vRNA copies/mL) for Zika virus/R.macaque-tc/UGA/1947/MR766-3329 (referred to as challenge stock) and ZIKV FP, respectively, were used for challenges. An additional isolate of ZIKV prototype strain MR766 with 150 suckling mouse brain passages and a single round of amplification on Vero cells was obtained from Robert Tesh (WRCEVA, Galveston, TX). After receipt, this virus also was amplified on C6/36 cells to produce stock virus (referred to as WRCEVA). See Table 1 for a full description of MR766 viruses.

**Primary challenge**. ZIKV MR766 challenge stock was thawed, diluted with PBS to the appropriate concentration for each challenge, and loaded into a 1mL syringe maintained on ice until challenge. For primary challenges, each of three, Indian-origin rhesus macaques was anesthetized and inoculated with 1mL subcutaneously over the cranial dorsum with either 1×10^4^, 1×10^5^, or 1×10^6^ PFU/mL of challenge stock. All animals were closely monitored by veterinary and animal care staff for adverse reactions and signs of disease. Animals were examined, and blood, pan urine, and oral swabs were collected from each animal daily from one through ten days post inoculation (dpi) and then weekly thereafter through 28 dpi. After 28 dpi, animals were rested for six weeks prior to secondary/heterologous challenge. Baseline sampling prior to secondary challenge occurred 56, 63, and 67 days post primary challenge.

**Secondary/heterologous challenge**. Seventy days after primary challenge, ZIKV FP was thawed and diluted with PBS to 1 × 10^4^ PFU/mL, loaded into a 1mL syringe and maintained on ice until challenge. Each animal was anesthetized and inoculated with 1mL subcutaneously over the cranial dorsum with 1×10^4^ PFU/mL ZIKV FP Animals were closely monitored by veterinary and animal care staff for adverse reactions and signs of disease. As described previously, animals were examined, and blood, urine, and saliva were collected from each animal daily from one through ten dpi and then weekly thereafter through 28 dpi.

**Plaque reduction neutralization test (PRNT90)**. Macaque serum samples were screened for ZIKV neutralizing antibody utilizing a plaque reduction neutralization test (PRNT). Endpoint titrations of reactive sera, utilizing a 90% cutoff (PRNT90) were performed as described (Lindsey et al., 1976) against ZIKV FP and challenge stock MR766.

**Viral RNA isolation**. Plasma was isolated from EDTA-anticoagulated whole blood collected the same day by Ficoll density centrifugation at 1860 rcf for 30 minutes. Plasma was removed to a clean 15mL conical tube and centrifuged at 670 rcf for an additional eight minutes to remove residual cells. Urine was opportunistically collected from a pan beneath each animal’s cage and centrifuged at 500 rcf for five minutes to remove cells and debris. Saliva was collected using sterile oral swabs run under the tongue while animals were anesthetized. Swabs were placed in viral transport media (tissue culture medium 199 supplemented with 0.5% FBS and 1% antibiotic/antimycotic) for 60-90 minutes, then vortexed vigorously and centrifuged at 500 rcf for five minutes. Prior to extraction, swab samples were pelleted by centrifugation at 14000 rpm and 4°C for an hour. After centrifugation, supernatant was removed, leaving virus in 200 μL media. Viral RNA was extracted from 300 μL plasma or urine using the Viral Total Nucleic Acid Kit (Promega, Madison, WI) on a Maxwell 16 MDx instrument (Promega, Madison, WI). Viral RNA was extracted from 200 μL oral swab-derived samples using the QIAamp MinElute Virus Spin Kit (Qiagen, Germantown, MD) with all optional washes. RNA was then quantified using quantitative RT-PCR. Viral load data from plasma and urine are expressed as vRNA copies/mL. Viral load data from oral swabs are expressed as vRNA copies/mL eluate.

**Quantitative reverse transcription PCR (qRT-PCR)**. For both ZIKV MR766 and ZIKV FP, vRNA from plasma, urine, and oral swabs was quantified by qRT-PCR using primers with a slight modification to those described by Lanciotti et al. to accommodate the anticipated African ZIKV sequences (Lanciotti et al., 2008). The modified primer sequences are: forward 5’-CGYTGCCCAACACAAGG-3’, reverse 5’-CACYAAYGTTCTTTTGCABACAT-3’, and probe 5’-6fam-AGCCTACCTTGAYAAGCARTCAGACACYCAA-BHQ1-3’. The RT-PCR was performed using the SuperScript III Platinum One-Step Quantitative RT-PCR system (Invitrogen, Carlsbad, CA) on a LightCycler 480 instrument (Roche Diagnostics, Indianapolis, IN). The primers and probe were used at final concentrations of 600 nm and 100 nm respectively, along with 150 ng random primers (Promega, Madison, WI). Cycling conditions were as follows: 37°C for 15 min, 50°C for 30 min and 95°C for 2 min, followed by 50 cycles of 95°C for 15 sec and 60°C for 1 min. Viral RNA concentration was determined by interpolation onto an internal standard curve composed of seven 10-fold serial dilutions of a synthetic ZIKV RNA fragment based on ZIKV FP. A comparison of the crossing point detected by qRT-PCR from the standard template, ZIKV FP and ZIKV MR766 when using the universal primer set developed by our group suggests that efficiency of these primers is the same for both lineages of ZIKV and comparable to the efficiency of those designed by Lanciotti et al. for Asian ZIKV (Supplemental Figure 1).

**Deep sequencing**. A vial of the same ZIKV MR766 stock used for primary challenge (i.e., challenge stock), a vial of the CDC MR766 stock, and a vial of the WRCEVA MR766 stock were each deep sequenced by preparing libraries of fragmented double-stranded cDNA using methods similar to those previously described (Lauck et al., 2013). Briefly, the sample was centrifuged at 5000 rcf for five minutes. The supernatant was then filtered through a 0.45-μm filter. Viral RNA was isolated using the QIAamp MinElute Virus Spin Kit (Qiagen, Germantown, MD), omitting carrier RNA. Eluted vRNA was then treated with DNAse I. Double-stranded DNA was prepared with the Superscript Double-Stranded cDNA Synthesis kit (Invitrogen, Carlsbad, CA) and priming with random hexamers. Agencourt Ampure XP beads (Beckman Coulter, Indianapolis, IN) were used to purify double-stranded DNA. The purified DNA was fragmented with the Nextera XT kit (Illumina, Madison, WI), tagged with Illumina-compatible primers, and then purified with Agencourt Ampure XP beads. Purified libraries were then sequenced with 2 × 300 bp kits on an Illumina MiSeq.

Virus populations replicating in plasma were sequenced using methods similar to those described previously (Gellerup et al., 2016). Viral RNA was isolated from 500 μl of plasma using the QIAamp MinElute Viral RNA isolation kit, according to manufacturer’s protocol. Viral RNA was then subjected to RT-PCR using the Superscript III One-step RT-PCR kit (Invitrogen, Carlsbad, CA), MgSO4, and 1.2uM of the primer pairs ZUG-1F: TCAACAGATGGGGTTCCGTG; ZUG-1R: GGGGGAGTCAGGATGGTACT. The following cycling conditions were used: 55°C for 30 min; 94°C 2 min; 35 cycles of the following: 94°C 15 sec, 56°C 30 sec, and 68°C 3.5 min; 68°C 10 min. Viral cDNA amplicons were size selected by agarose gel electrophoresis and then purified using the Qiagen MinElute Gel Extraction kit. Purified PCR products were pooled and then ~1 ng of DNA was fragmented using the Nextera XT kit (Illumina), tagged with Illumina-compatible primers, and then purified with Agencourt Ampure XP beads. Purified libraries were then sequenced with 2 × 300 bp kits on an Illumina MiSeq.

Sequences were analyzed using a modified version of the viral-ngs workflow developed by the Broad Institute (http://viral-ngs.readthedocs.io/en/latest/description.html) and implemented in DNANexus. Briefly, host-derived reads that map to a human sequence database and putative PCR duplicates are removed. The remaining reads were mapped to an NCBI Genbank MR766 reference sequence (HQ234498). The published viral-ngs workflow uses the Novoalign read mapper; however, Novoalign is relatively insensitive to the 12 nucleotide in-frame deletion in the MR766 envelope. Therefore, we modified the viral-ngs pipeline to use the bwa mem version 1.5.0 (http://bio-bwa.sourceforge.net) read mapper with default parameters to map reads sequence reads to HQ234498. Deep sequencing datasets are available from http://go.wisc.edu/50bfn2 and are deposited in the NCBI Sequence Read Archive with accession numbers (in process); the DNANexus workflow for read mapping is available upon request from the authors.

Mapped reads and reference scaffolds were loaded into Geneious Pro (Biomatters, Ltd., Auckland, New Zealand) for intrasample variant calling. Variants were called in Envelope that fit the following conditions: present in >=5% of the mapped reads, have a minimum p-value of 10e-60, a minimum strand bias of 10e-5 when exceeding 65% bias, and were nonsynonymous. Variant call format files are available from http://go.wisc.edu/50bfn2. Mapping metrics can be found in Supplemental Figure 2.

**Comparison of East African ZIKV MR766 and Asian Zika virus isolates**. Full-length Asian-lineage Zika virus sequences available in NCBI Genbank as of June 8, 2016 were copied into Geneious Pro 9.1.2 (Biomatters, Ltd., Auckland, New Zealand). The amino acid sequence of E was obtained from these sequences, as well as the consensus sequence from MR766 challenge stocks, by conceptual translation. These amino acid sequences were aligned with MUSCLE (Edgar, 2004) as implemented in Geneious Pro 9.1.2 using default parameters.

**Immunophenotyping**. Numbers of activated and proliferating NK cells were quantified as described previously (Pomplun et al., 2015). For each timepoint analyzed, 100 μL of EDTA-anticoagulated whole blood samples were incubated at room temperature for 15 minutes with an antibody master mix described in detail in (Dudley et al., 2016). Red blood cells were lysed (BD Pharm Lyse, BD Biosciences, San Jose, CA), washed twice, and then fixed with 2% paraformaldehyde for 15 minutes. After fixation, cells were washed and permeabilized using Bulk Permeabilization Reagent (Life Technologies, Madison, WI) and stained with Ki-67 (clone B56, Alexa Fluor 647 conjugate) for 15 minutes. After staining, cells were washed again and resuspended in 2% paraformaldehyde until use in flow cytometry (BD LSRII Flow Cytometer, BD Biosciences, San Jose, CA). Flow cytometry data were analysed using FlowJo v. 9.9.3 (TreeStar, Ashland, OR).

**PBMC processing**. Fresh PBMC were isolated by Ficoll gradient as described in vRNA isolation. PBMC were collected into R10 media (Hyclone, Logan, UT) and centrifuged at 670 rcf for five minutes, treated with ACK (Grand Island, NY) to removed residual RBC, washed twice more with R10 media, and centrifuged again. R10 was removed and cells were resuspended in Cryostor CS5 media (BioLife Solutions, Bothell, WA), and frozen (1°C/ minute) down to −80°C storing in liquid nitrogen vapor phase until plasmablast assays were performed.

**Plasmablast detection**. PBMCs isolated from the three ZIKV-002 macaques were stained with the following panel of fluorescently labeled antibodies (Abs) specific for the following surface markers: CD20 FITC (L27), CD80 PE(L307.4), CD123 PE-Cy7(7G3), CD3 APC-Cy7 (SP34-2), IgG BV605(G18-145) (all from BD Biosciences, San Jose, CA), CD14 AF700 (M5E2), CD11c BV421 (3.9), CD16 BV570 (3G8), CD27 BV650(O323) (all from BioLegend, San Diego, CA), IgD AF647 (polyclonal)(Southern Biotech, Birmingham, AL), and HLA-DR PE-TxRed (TU36) (Invitrogen, Carlsbad, CA). LIVE/DEAD Fixable Aqua Dead Cell Stain Kit (Invitrogen, Carlsbad, CA) was used to discriminate live cells. Briefly, cells were resuspended in 1X PBS/1%BSA and stained with the full panel of surface Abs for 30 min in the dark at 4°C, washed once with 1X PBS, stained for 30 min with LIVE/DEAD Fixable Aqua Dead Cell Stain Kit in the dark at 4°C, washed once with 1X PBS, washed again with 1X PBS/1%BSA, and resuspended in 2% PFA Solution. Stained PBMCs were acquired on a LSRII Flow Analyzer (BD Biosciences, San Jose, CA) and the data was analyzed using FlowJo software v9.7.6 (TreeStar, Ashland, OR). Plasmablasts were defined similarly to the method previously described (Silveira et al., 2015) excluding lineage cells (CD14+, CD16+, CD3+, CD20+, CD11c+, CD123+), and selecting CD80+ and HLA-DR+ cells (known to be expressed on rhesus plasmablasts and their human counterpart (Wrammert et al., 2008).

## Author contributions

D.H.O, M.T.A., D.M.D., T.C.F., S.L.O., E.L.M., J.E.O., and T.G.G. designed the experiments. D.H.O., M.T.A., D.M.D., and S.L.O. drafted the manuscript. All authors edited the manuscript. M.T.A. and J.E.O. provided and prepared viral stocks and performed PRNTs. M.E.B., C.R.B., M.S.M., M.N.R., C.M.N., and D.M.D. coordinated and processed macaque samples for distribution. A.M.W, G.L.B. and T.C.F. performed viral load assays. D.D.G., L.H.M, and S.L.O. designed and performed the sequencing experiments. M.N.R., K.L.W, L.J.V, and E.G.R performed immunophenotyping assays. J.P. and S.C. coordinated macaque infections and sampling. S.R.P., M.A.M, and J.A.E performed the plasmablast experiment.

## Acknowledgements

The authors acknowledge Jens Kuhn and Jiro Wada for preparing silhouettes of macaques used in figures. This project would not have been possible without the support of a supplement to NIH grant 1R01AI116382-01A1. Research reported in this publication was also supported in part by the Office Of The Director, National Institutes of Health under Award Number P51OD011106 to the Wisconsin National Primate Research Center, University of Wisconsin-Madison. This research was conducted at a facility constructed with support from Research Facilities Improvement Program grant numbers RR15459-01 and RR020141-01. The content is solely the responsibility of the authors and does not necessarily represent the official views of the National Institutes of Health.

## Supplemental information

**Figure S1.**
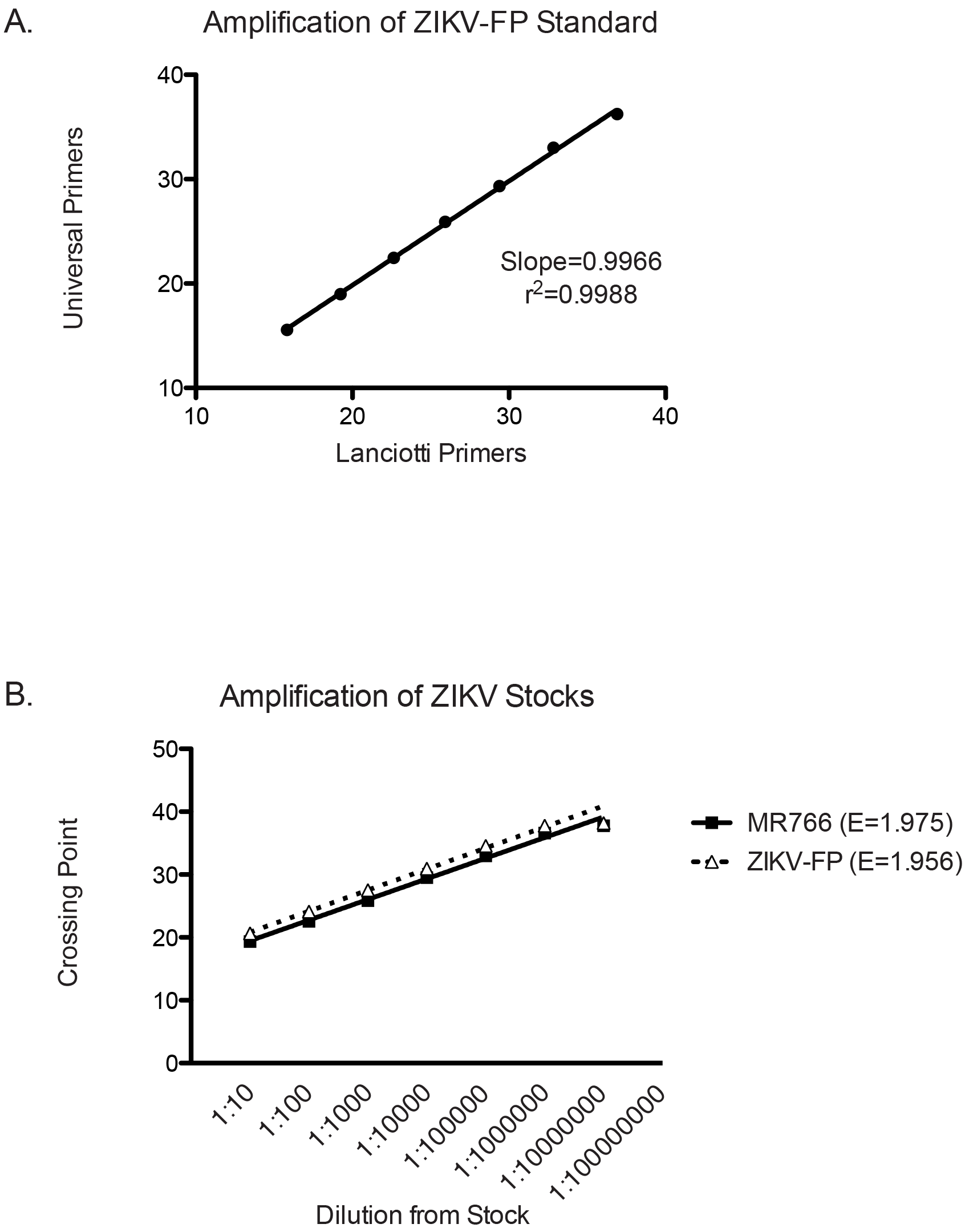
Validation of Universal Primers by qRT-PCR. Crossing point indicates threshold PCR cycle at which amplification was first detected. A. Comparison of crossing points seen in amplification of a synthetic ZIKV-FP standard curve using our universal primers and those designed by Lanciotti et al {18680646}. B. Comparison of amplification efficiencies of universal primers for East African MR766 and ZIKV-FP targets. Universal primers were used in qRT-PCR to amplify serial tenfold dilutions of MR766 or ZIKV-FP stocks. Amplification efficiencies were 1.975 and 1.956 for MR766 and ZIKV-FP, respectively, with 2 being a theoretically perfect efficiency (i.e., DNA concentrations double each cycle).

**Supplemental Figure 2:**
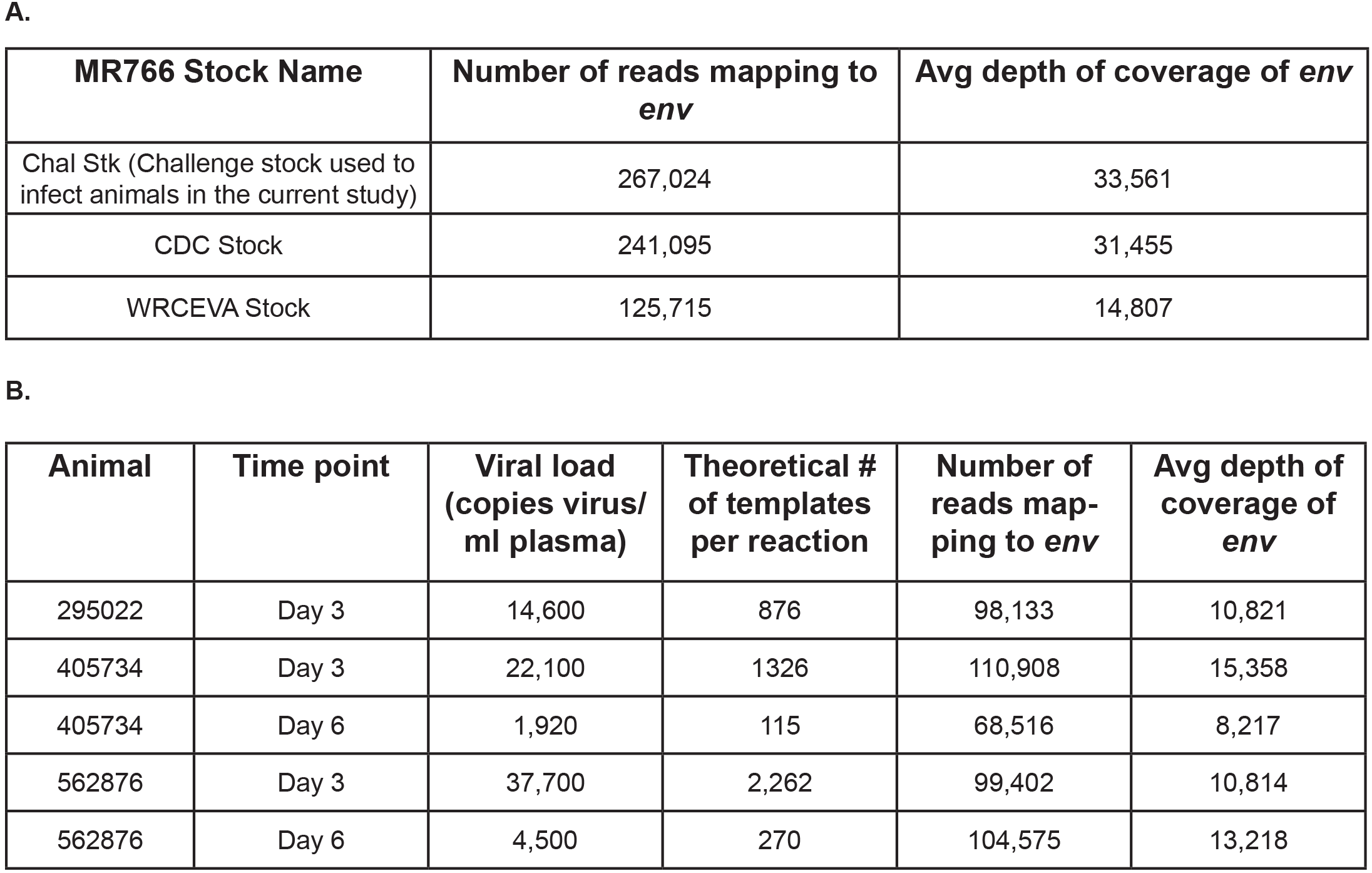
Metrics for deep sequencing of virus stocks and samples from animals. A. Metrics for the sequences of the three stocks (Figure 1A) spanning the env gene are shown. The number of individual reads mapping to env and the average depth of coverage are shown. B. Metrics for the env sequences generated from animals (Figure 3) are shown. Theoretical number of templates was calculated by assuming that isolation of viral RNA from 500ml of plasma was complete into 25ml of elution buffer, and was followed by using 3ml of eluted viral RNA per RT-PCR reaction. The number of individual reads mapping to env and the average depth of coverage are shown.

